# Antagonistic Regulation of LINE-1/Alu Elements and Their Repressor APOBEC3B in Cellular Senescence

**DOI:** 10.1101/2025.03.27.645653

**Authors:** Daksha Munot, Gregoire Najjar, Manvendra Singh, Cagatay Günes, Daniel Sauter, Ankit Arora

## Abstract

Long Interspersed Nuclear Elements-1 (LINE-1 or L1) make up approximately 21% of the human genome, with some L1 loci containing intact open reading frames (ORFs) that facilitate retrotransposition. Because retrotransposition can have deleterious effects leading to mutations and genomic instability, L1 activity is typically suppressed in somatic cells through transcriptional and post-transcriptional mechanisms. However, L1 elements are derepressed in senescent cells causing age-associated inflammation. Despite the recognition of L1 activity as a hallmark of aging, the underlying molecular mechanisms governing L1 derepression in these cells are not fully understood. In this study, we employed high throughput sequencing datasets and validated our findings through independent experiments to investigate the regulation of L1 elements in senescent cells. Our results reveal that both replicative and oncogene-induced senescence are associated with reduced expression of the cytidine deaminase *APOBEC3B*, a known suppressor of L1 retrotransposition. Consequently, senescent cells exhibited diminished levels of C-to-U editing of full-length L1 elements. Moreover, Ribo-seq profiling indicated that progression to senescence is not only associated with increased L1 transcription, but also translation of L1 ORFs. In summary, our results suggest that the depletion of APOBEC3B contributes to enhanced activity of L1 in senescent cells and promotion of L1-induced DNA damage and aging.

## Background

Almost half of the human genome consists of sequences derived from transposable elements (TEs) [1]). The majority of TE sequences belong to non-LTR retrotransposon families such as L1 (21% of the genome with ∼ 500,000 copies) and Alu elements (∼ 10% of the genome with ∼ 1 million copies) [2,3]. While most L1 elements are inactive, a few full-length loci are still capable of retrotransposition [1]. Moreover, retrotransposon families that do not encode their own retrotransposition machinery (e.g. Alu and SVA elements) also exploit L1-encoded reverse transcriptase to integrate into new genomic loci [4,5].

Due to the potential harmful effects of active TEs, such as insertional mutagenesis, the human genome has evolved independent mechanisms to suppress TEs at multiple steps of their retrotransposition cycle. For instance, specific KRAB-ZNFs recruit TRIM28 (Tripartite motif-containing protein 28) that silences L1 transcription elements by further recruiting chromatin-modifying enzymes such as the histone methyltransferase SETDB1 [6]. SETDB1 is also known to interact with the HUSH complex (TASOR, Periphilin, MPP8), causing long-term repression by perpetuating heterochromatin [7]. However, even if L1 elements are transcriptionally active, their retrotransposition can still be curbed at later steps. For instance, nucleases such as three prime repair exonuclease 1 (TREX1) prevent the accumulation of L1 transcripts [8,9], while MOV10 decaps L1 RNA [10], thereby sequestering L1 ribonucleoprotein complexes in cytoplasmic aggregates [11]. A critical inhibition of retrotransposition is mediated by the members of the APOBEC (Apolipoprotein B mRNA Editing Catalytic Polypeptide-like) family of deaminases [12–20]. While APOBEC3 proteins are known to induce C-to-U mutations, leading to hypermutation, their role in suppressing L1 retrotransposition appears to extend beyond this deaminase activity [4]. Notably, overexpression of APOBEC3A, B, C, and D inhibits L1 activity without corresponding increases in L1 point mutations, indicating a deaminase-independent mechanism [14]. Further supporting this, catalytically inactive mutants of APOBEC3B and C retain their ability to suppress L1 retrotransposition underscoring the multifaceted roles these proteins play in maintaining genomic stability [15,20].

Despite these multilayered suppression mechanisms, L1 elements can be activated under certain conditions. For example, external stimuli such as ionizing radiation trigger L1 transcription [21]. Similarly, certain developmental stages are associated with increased L1 activity. These include early developmental stages such as embryonic cells that undergo epigenetic reprogramming and global demethylation [22]. Notably, DNA methylation of L1 repeats decreases with advancing age, and L1 activity has been proposed as a predictor of chronological age [23,24]. In this case, L1 activity was shown to drive organismal aging and cellular senescence [25,26]. Senescence describes a state of permanent cell-cycle arrest and resistance to apoptosis that can be induced by multiple intrinsic and extrinsic stimuli, including genotoxic stress or oncogene activation [27]. It is typically associated with the release of inflammatory cytokines and other immune modulators, referred to as the senescence-associated secretory phenotype (SASP) [28]. These secreted factors can trigger the progression of proliferating, non-senescent cells to senescence [28].

The activation of L1 repeats in pre-senescent cells results in the production of nucleic acids that can activate cellular sensing pathways such as cGAS-STING or RIG-I, which mediate the production of type I IFN and ultimately SASP [25,26,29]. Hence, L1 activity in pre-senescent cells not only contributes to their transition to senescence, but can also induce a senescent state in bystander cells [30]. In addition to sensing of L1-derived nucleic acids, the retrotransposition and *de-novo* integration of L1 elements may promote senescence by triggering the vicious cycle of DNA damage, cell-cycle arrest and inflammation [26]. Overall, there is more than one way how uncontrolled activity of L1 can trigger cellular senescence.

While SASP is heterogenous, and numerous triggers of senescence have been described, L1 transcription has been observed in different types of senescence, including replicative senescence, oncogene-induced senescence and stress-induced premature senescence [25,31]. L1 activation in senescent cells is associated with higher levels of accessible chromatin around the 5’ UTR of L1 repeats [31]. Furthermore, the transcription factor PAX5 binds to the 5’-UTR of L1 and is proposed to contribute to their activation in cells entering senescence [32]. However, precise mechanism of enhanced L1 activity in (pre)senescent cells remains elusive, and the mechanisms underlying post-transcriptional L1 regulation are still unclear.

We hypothesized that the loss of various L1 controllers may result in L1 derepression at the transcriptional and/or post-transcriptional level. To investigate the expression dynamics of L1 and its repressors, we reanalyzed publicly available RNA-seq and Ribo-seq datasets from cells progressing to replicative or oncogene-induced senescence.

Our RNA-seq data analyses, together with validation experiments in fibroblasts, revealed that several L1 inhibitors, including the deaminase APOBEC3B, are down-regulated in presenescent cells. In line with reduced APOBEC3B levels, we found evidence for decreased C-to-U editing and higher levels of L1 translation. In summary, our findings support a model in which reduced expression of restriction factors such as APOBEC3B contribute to increased L1 activity, ultimately leading to enhanced L1 cDNA sensing and potentially L1-mediated DNA damage in cells transitioning to senescence.

## Materials and Methods

### Analysis of RNA-seq datasets from senescent cells

Raw RNA-seq reads from immortalized human primary BJ fibroblasts and ES-derived lung fibroblasts were obtained from [33] [GSE42509] and [25] [GSE109700], respectively. RNA-seq reads were mapped to the human (hg19) reference genome using STAR (v2.4.2a) [34]. We used STAR for its ability to detect novel splice junctions and as a non-canonical splice aligner for the detection of chimeric transcripts and circular RNA. Mapped reads were processed to obtain raw counts for gene expression estimation using featureCounts [35] from the subread package (v1.4.6). DESeq2 (v1.30.1) was used to process counts per million (CPM) and differential expression of genes in the R environment (v4.0.3). To determine the expression of cellular factors affecting L1 retrotransposition, we took advantage of a list of activators and suppressors of L1 retrotransposition previously identified in a genome-wide CRISPR/Cas9 screening [36], as well as experimentally validated and published L1 suppressors (Additional files 1 and 2).

### Analysis of repetitive elements

Mapping of RNA-seq reads to the human reference genome (hg19) was performed using bowtie (v1.0.1) [37] short read aligner with use of -N 1 and - -local parameters for effective alignment. FPKM values for expression were computed using cufflinks (v2.0.2) [38] with the usage of GTF file format for specific repetitive elements (L1 & Alu) obtained from the UCSC genome browser [39]. The annotation file of repetitive elements (L1 & Alu) was categorized into young, middle-aged and ancient sub-families.

### Detection of APOBEC- and ADAR-edited RNA sites

A data analysis pipeline was designed, involving multiple rigorous iterations focused on the identification of APOBEC- and ADAR-edited RNA sites within full-length L1 elements. Mapped RNA-seq reads were incorporated to perform local realignment, base-score recalibration, and candidate variant calling using the IndelRealigner, TableRecalibration and UnifiedGenotyper tools with the parameters stand call conf to 0 and stand emit conf to 0 and the output mode set to EMIT ALL CONFIRMED SITES from the Genome Analysis Toolkit (GATK) (v3.5-0) [36,40]. In addition, some iteration steps of the SNPiR pipeline [41] were adapted, and obtained variants were subjected to it. The intended variants were incorporated into different filtering steps to obtain true variants by removing false-positive variant calls. First, variants with quality up to 20 were filtered. Then, the mismatches at the 5’ ends of the reads were removed in this step. Furthermore, the obtained variants were directed to filter in L1 elements. Shell scripting and bedtools [42] were used to retrieve the edited sites for each sample across full-length L1 elements. The obtained editing sites for each sample were further filtered for C to U mutations for APOBEC editing sites and A to I mutations for ADAR editing sites.

### Estimation of L1 encoded proteins

The repeat masker track for L1 elements was downloaded from the UCSC genome browser. More specifically, only full-length L1 elements were extracted from this track, and a BED file was prepared. Ribosomal profiling (Ribo-seq) samples for immortalized human primary BJ fibroblasts were obtained from [33] [GSE42509]. Adapter sequences were removed from the raw FASTQ reads, and ribosomal sequences were eliminated using quick alignment with TopHat2 [43]. Unmapped reads from this alignment were converted into FASTQ format using bam2fastx [43]. The consensus sequence for full-length L1 elements was constructed by converting the BED file into FASTA format with fastaFromBed [42]. These consensus sequences were aligned to Ribo-seq reads (single-end, 50 bp) using Bowtie (v1.0.1) using the best parameter, across conditions of proliferation and senescence. Uniqueness for 35-mers and alignability for 36-mers with respect to full-length L1 elements in the Ribo-seq data were computed. Normalized coverage for uniquely mapped reads was obtained using bamCoverage for each condition. Further, computation of matrix was performed on the bigwig file of each condition by using computeMatrix [44].

### Cell culture, transfection and transduction

BJ cells were purchased from ATCC (ATCC LGC Standards GmbH, Manassa, Virginia, USA; cat# CRL-2522). BJ-hTERT cells were generated by transduction with the pWZL-Blast-Flag-HA-hTERT retroviral vector (addgene; cat.# 22396). All cells were cultured in DMEM high glucose, penicillin (10.000 unit/mL), streptomycin (10 mg/mL), and 10% FCS at 37 °C and 5% CO_2_. HEK293T cells were transfected with PWZL-Hygro-H-RAS^G12V^ using PEI. On the following day, the cell culture medium was changed and 24 hours later a supernatant containing the viral particles was filtered through a 0.45 µm filter. Viral supernatant and 8 µg/µL polybrene were then added to the target cells (primary BJ and immortalized BJ-hTERT) for 24 hours. Lastly, cells were washed three times with DPBS and appropriate selection antibiotic was added.

### qRT-PCR

At 7 days post-transduction (d.p.t.), total RNA was isolated using the RNeasy Mini Kit (Qiagen, cat# 74106) following the manufacturer’s instructions. RNA quality and quantity were assessed using a spectrophotometer. Reverse transcription was performed using the PrimeScript RT Reagent Kit (Perfect Real Time) (TAKARA, cat# RR037A) with oligo dT primers and random hexamers. Quantitative real-time PCR (qPCR) was conducted using specific primer/probe sets for *APOBEC3B* (Thermo Fisher, cat# Hs00358981_m1), with *GAPDH* (Thermo Fisher, cat# Hs02786624_g1) serving as the internal control. Each sample was analyzed in technical triplicates.

### Statistical analysis

Wilcoxon-Mann-Whitney test, paired t-test and Ordinary one-way ANOVA test were used for analysis at the transcriptional and translational level. p-values <0.05 were considered statistically significant.

## Results

### Replicative and oncogene-induced senescence are associated with decreased expression L1 repressors such as *APOBEC3B*

While L1 activity was shown to be upregulated in senescence [25,31], the mechanisms underlying L1 activation remained poorly understood. We therefore commenced this study by determining the expression levels of known L1 suppressors in primary human lung fibroblasts transitioning from a proliferative state to early and ultimately late replicative senescence. By leveraging a previously published RNA-seq dataset [25] [GSE109700], we compared the expression of L1 suppressors in proliferative cells relative to both early and late senescent cells taken together. We found 13 L1 repressors to be significantly downregulated in senescent cells compared to proliferating cells (Fig. 1A, Additional file 1). The five most strongly reduced transcripts encode for the deaminase APOBEC3B (log2fc -5.79 & adjusted p-value 1.28e-04) and the DNA repair proteins BRCA2 (log2fc -4.75 & adjusted p-value 3.56e-36), FANCD2 (log2fc -4.58 & adjusted p-value 1.51e-38), FANCA (log2fc -4.57 & adjusted p-value 3.27e-83) and BRCA1 (log2fc -4.51 & adjusted p-value 2.98e-127) (Additional file 1). In contrast, only a single L1 suppressor, SLFN5, was significantly upregulated during progression to senescence (log2fc 1.45 & adjusted p-value 2.17e-15) (Additional file 1). The data suggest that progression to senescence is marked by a significant downregulation of multiple L1 suppressors, particularly APOBEC3B, BRCA2, FANCD2, FANCA, and BRCA1, while SLFN5 is the only L1 repressor significantly upregulated.

**Fig. 1:**
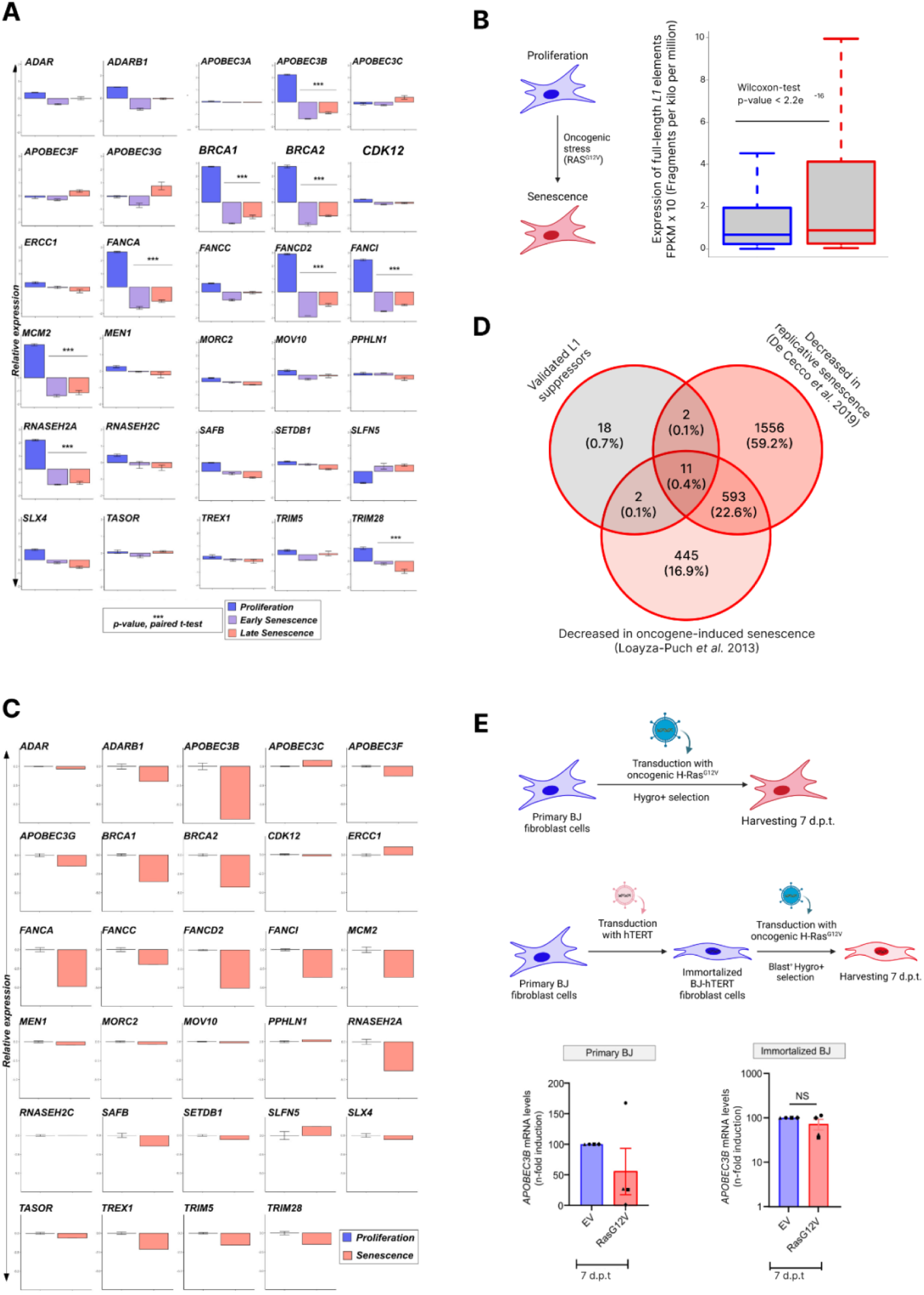
Decreased expression of L1 repressors in senescent cells. **(A)** Expression of known L1 suppressors in human lung fibroblasts transitioning from a proliferative state to early and late replicative senescence (see also Additional file 1). Expression relative to the mean of all three conditions is shown. A paired t-test was performed to determine statistically significant differences between proliferating and senescent (early+late) cells (***p<0.001). **(B)** Expression of full-length L1 elements is significantly increased in senescent vs. proliferating cells (p-value < 2.2e-16). (**C**) Expression of known L1 suppressors in BJ fibroblasts before and after RAS^G12V^-mediated induction senescence. The analyses in (A, B, and C) are based on publicly available RNA-seq datasets (GSE109700 and GSE42509, respectively). **(D)** Venn diagram illustrating the overlap of known L1 suppressors with genes that are down-modulated during transition to replicative and/or oncogene-induced senescence. **(E)** Cartoon illustrating the transduction process of hTERT-negative and hTERT-immortalized BJ fibroblast cells is shown on top. *APOBEC3B* mRNA levels upon transduction with lentiviruses encoding the oncogene RAS^G12V^ or the respective control (EV) are shown at the bottom. Mean values of four independent experiments ± SEM are shown (*p<0.05; NS not significant, d.p.t. days post-transduction)

To determine whether a similar expression pattern of L1 inhibitors can also be observed in other models of senescence, we determined their expression in cells undergoing oncogene-induced senescence [25]. To this end, we re-analyzed RNA-seq data obtained from immortalized human BJ primary fibroblasts [GSE42509], in which senescence was induced by tamoxifen-inducible expression of the oncogenic *RAS^G12V^* gene [33]. Since this study did not monitor the activity of transposable elements, we first analyzed the transcription of L1 elements in this data set. As observed for replicative senescence, L1 expression in BJ fibroblasts was significantly elevated upon *RAS^G12V^*-mediated induction of oncogenic senescence (Fig. 1B). Furthermore, oncogene-induced senescence was also associated with a significant decrease in the expression of L1 inhibitors (Fig. 1C, Additional file 2). Intriguingly, eleven L1 repressors were significantly depleted in both data sets (Additional files 1 and 2, Fig. 1D). In oncogene-induced senescence, the five most strongly depleted transcripts were *APOBEC3B* (log2FC -6.75 & adjusted p-value 4.08e-11), *FANCD2* (log2FC -4.84 & adjusted p-value 1.51e-38), *FANCA* (log2FC -4.68 & adjusted p-value 6.35e-21), *BRCA2* (log2FC -4.02 & adjusted p-value 4.09e-18) and *RNASEH2A* (log2FC -3.62 & adjusted p-value 8.21e-12) (Fig. 1C, Additional file 2). Again, expression of *SLFN5* was modestly increased in senescent vs. proliferating cells (log2FC 1.20 & adjusted p-value 0.43). These findings indicate that both replicative and oncogene-induced senescence are marked by a significant downregulation of multiple L1 inhibitors.

Next, we expanded our analyses to a list of potential L1 regulators identified by Liu and colleagues [36]. While the activity of many of the identified activators and inhibitors remains to be experimentally validated, the candidate L1 regulators are the results of unbiased genome-wide CRISPR screens [36]. Several L1 suppressors were down-regulated in cells progressing from proliferation to senescence (Additional files 3 and 4). This was particularly evident for cells undergoing replicative senescence (Additional file 3B). Surprisingly, several L1 activators also showed reduced expression in cells transitioning to (replicative) senescence (Additional file 3C, D).

Since *APOBEC3B* was the most strongly down-regulated repressor in both replicative and oncogene-induced senescence (Fig. 1A, C, Additional files 1, and 2), we validated its differential expression via qRT-PCR. To this end, we took advantage of the RAS^G12V^ fibroblast model, including telomerase reverse transcriptase (hTERT) positive and negative cells. As previously described [94,95], hTERT-negative BJ cells show signatures of premature senescence 7 days post transduction with RAS^G12V^. In contrast, hTERT-immortalized BJ fibroblasts do not enter senescence, but stay in a growth-delayed state between 7-14 days after Ras^G12V^ expression that is ultimately overcome [94,95] (Fig. 1E, top). 7 days after Ras^G12V^ expression, *APOBEC3B* levels decreased 1.8-fold in presenescent, telomerase-negative cells. In contrast, a 1.4-fold reduction of *APOBEC3B* expression was observed in hTERT-expressing cells (Fig. 1E, bottom). Together, these findings demonstrate that transition of proliferating cells to oncogene-induced and replicative (pre)senescence is associated with reduced expression of the restriction factors *APOBEC3B* and a variety of other L1 inhibitors.

### Expression of Alu element subfamilies is increased in senescent cells

Many L1 repressors also restrict other transposable elements. For example, APOBEC3 family members, including APOBEC3B [16,19,27,45–47], also suppress Alu retrotransposition, and a subset of Alu repeats is bound by TRIM28 [27,45]. We therefore expanded our analyses to different Alu elements, which belong to the group of short interspersed nuclear elements (SINEs). These include the two major Alu subfamilies AluJ and AluS [48], as well as AluY, a sub-subfamily of AluS [49]. Expression of all three Alu subfamilies was higher in oncogene-induced senescent cells compared to proliferating cells (p-values < 2.2e-16) (Additional file 4). These results demonstrate that reduced expression of TE suppressors is not only associated with increased L1 activity, but also with increased expression of the major Alu subfamilies.

### Senescent cells show reduced signatures of APOBEC-induced L1 RNA editing

APOBEC3B, the most strongly downregulated repressor in senescent vs. proliferating cells (Fig. 1A, C, Additional files 1 and 2), is able to deaminate both RNA and DNA [50]. Although APOBEC3 proteins are able to restrict L1 independently of their deaminase activity [14,15,20], we therefore hypothesized that the reduction of *APOBEC3B* expression may coincide with decreased RNA editing in cells transitioning to senescence. To test this hypothesis, we examined APOBEC3-mediated RNA editing profiles in cells progressing to oncogene-induced senescence [33]. We employed RNA-seq datasets for identification of RNA editing profiles. We incorporated mapped RNA-seq reads to execute local realignment, base-score recalibaration and candidate variant calling by using GATK toolkit. The obtained variants were incorporated to into various filtering steps to obtain true variants by removal of false-positive variant calls. Furthermore, the obtained variants were directed to filter in L1 elements. Using an in-house pipeline, we retrieved edited sites for each sample across full-length L1 elements (see Methods). When investigating the loci of ancient (e.g. L1PA6, L1PA7) and middle-aged L1 subfamilies (e.g. L1PA2, L1PA3, L1PA4, L1PA5), which are not capable of retrotransposition, we observed no significant difference in the editing frequency between proliferating and senescent cells (Fig. 2A). In contrast, the evolutionary youngest and most active L1 subfamilies (e.g. L1Hs, L1PA1) showed an ∼8 fold reduced editing frequency in senescent vs. non-senescent cells (Fig. 2B) (p-value = 0.024). As additional control, we also monitored editing signatures induced by ADAR (Adenosine Deaminase Acting on RNA) proteins, which convert adenosine into inosine [51]. ADAR1 and ADAR2 (=ADARB1) both restrict L1 [52,53], and ADARB1 was significantly downregulated in senescent vs. non-senescent cells (Additional files 1 and 2). However, we found no significant changes in ADAR-mediated editing signatures within L1 elements (Fig. 2C, D). In summary, our RNA editing analyses point towards a reduced mutagenic activity of at least one APOBEC3 protein in senescent cells.

**Fig. 2:**
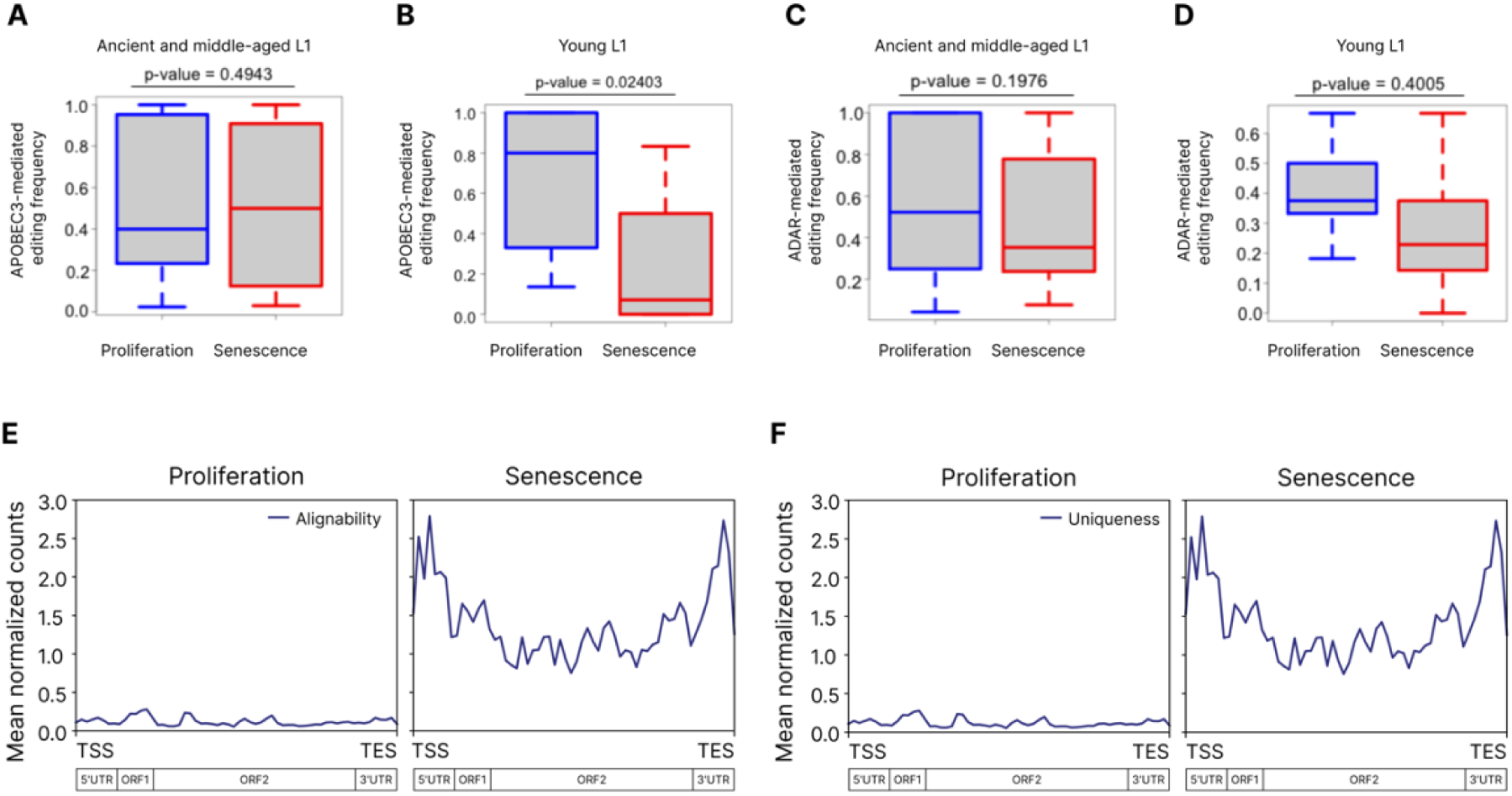
Editing and translation of L1 RNA in senescent vs. proliferating cells. **(A)** Frequency of APOBEC3-mediated editing of ancient (e.g. L1PA6, L1PA7) and middle-aged (e.g. L1PA2, L1PA3, L1PA4, L1PA5) L1 subfamilies. **(B)** Frequency of APOBEC3-mediated editing of the youngest L1 loci (e.g. L1HS), which are still capable of retrotransposition (p-value=0.02403). **(C)** Frequency of ADAR-mediated editing of ancient (e.g. L1PA6, L1PA7) and middle-aged (e.g. L1PA2, L1PA3, L1PA4, L1PA5) L1 subfamilies. **(D)** Frequency of ADAR-mediated editing of the youngest L1 loci (e.g. L1HS). Statistically significant differences in editing frequency were calculated using the Wilcoxon-Mann-Whitney test. The analyses in (A-D) are based on publicly available RNA-seq datasets (GSE42509. **(E) & (F)** Coverage-plot demonstrating the enrichment of Ribo-seq reads within L1 in senescence vs. proliferation. While (E) describes the enrichment for alignability, (F) describes the enrichment for uniqueness. Analyses in (E) and (F) are based on Ribo-seq datasets obtained from [33].

### Translation of L1 ORFs is increased in senescent cells

Increased L1 transcription and decreased L1 RNA editing in senescent cells are predicted to result in increased translation of L1 ORFs. We therefore examined publicly available Ribo-seq datasets of immortalized human primary fibroblasts in proliferative and senescent (oncogene-induced) states [33]. Due to the short sequencing reads (29-35 bps), the analysis of Ribo-seq datasets for TEs, especially the youngest ones, poses major alignability and mappability challenges [54]. To overcome these challenges, we aligned the Ribo-seq datasets with the full length L1 consensus sequence using an in-house pipeline and compared it to the genome-wide 35 bps unique mappability/alignability of full-length L1 elements. After calculating Ribo-seq coverage (see Methods), we were able to identify a few hundred mappable reads over the full-length L1 consensus sequence comprising both ORFs with a similar pattern of 35 bps mappability over different L1 elements (Fig. 2E). As expected, stronger signals were observed around the translation start sites (TSS) and translation end sites (TES), where ribosomes briefly pause at the start and stop codons, respectively (Fig. 2F). Taken together, the results demonstrate that progression to senescence is not only associated with increased transcription, but also increased translation of L1 elements.

## Discussion

Our study provides insights into the expression dynamics of TE regulators in presenescent and senescent cells that may enable the derepression of L1 and/or Alu elements and ultimately contribute to an irreversible cell cycle arrest and inflammaging. By analyzing publicly available RNA-seq datasets, we reveal a consistent decrease in the expression of various TE inhibitors in two major subtypes of senescence, i.e. replicative and oncogene-induced senescence. Vice versa, several TE activators are upregulated in (pre)senescent cells. These expression changes of TE modulators are tightly coupled with increased transcription and/or translation of L1 and Alu elements. Thus, our findings suggest that a major dysregulation of TE-regulating factors contributes to L1 and Alu derepression in cells transitioning from a proliferating to a senescent state.

Among the commonly down-regulated inhibitors were transcriptional repressors such as TRIM28/KAP1, DNA replication regulators such as MCM2, but also several RNAses (e.g. RNASEH2A) and DNA repair proteins (e.g. FANCA, FANCD2, FANCC, FANCI, BRCA1, BRCA2) (Fig. 1A, C, Additional files 1 and 2). The most strongly down-regulated L1 repressor was APOBEC3B. This deaminase is well known for its ability to restrict exogenous retroviruses such as HIV by inducing lethal hypermutations in the viral genome [55]. While it also restricts L1 elements, this inhibitory activity is independent of RNA editing [56]. Still, we observed signatures of APOBEC3-mediated editing of L1 transcripts. APOBEC3B was the only APOBEC3 family member significantly downregulated in oncogene-induced and replicative senescence (Additional files 1 and 2), strongly suggesting that it was responsible for the observed L1 editing and suggesting a shared regulatory mechanism which in turn driving L1 activation in senescence. One interesting aspect in this context is that in contrast to other *APOBEC3* genes, *APOBEC3B* is not IFN-inducible [57]. Thus, its expression is not expected to be increased upon L1-mediated IFN production.

Our analysis of RNA editing profiles in cells progressing to oncogene-induced senescence revealed distinct patterns among L1 subfamilies. Specifically, the evolutionarily youngest and most active L1 subfamilies, such as L1HS and L1PA1, exhibited approximately a ∼8 fold reduction in editing frequency in senescent cells compared to non-senescent cells (Fig. 2B). In contrast, no significant differences in editing frequency were observed at loci corresponding to ancient (e.g., L1PA6, L1PA7) and middle-aged L1 subfamilies (e.g., L1PA2, L1PA3, L1PA4, L1PA5), which are no longer capable of retrotransposition. These findings suggest that the reduced RNA editing activity in senescent cells primarily affects the youngest and most active L1 elements, potentially limiting their mutagenic potential during senescence. This observation aligns with our hypothesis that the activity of APOBEC3 proteins, which are key mediators of L1 RNA editing, is diminished in (pre)senescent cells, thereby contributing to a reduced mutagenic burden in this state.

The reduction of various host factors that inhibit different steps of L1 retrotransposition suggests that L1 elements are not only transcribed, but also translated when cells progress to a senescent state [58]. Indeed, ribosomal profiling analyses demonstrate that L1 ORFs are translated in senescent cells. As we only analyzed retrotransposition-competent copies of full-length L1 elements, the above results suggest that senescent cells are characterized by increased ‘jumping’ of L1.

Since Alu elements depend on L1 for their transposition and since L1 inhibitors frequently also restrict Alu repeats [59,60,61,62], it is not surprising that expression of the latter is also increased in senescent vs. proliferating cells. Notably, this was not only the case for the youngest and most active sub-subfamily AluY, but also the oldest subfamily AluJ.

Since TEs of (pre)senescent cells are derepressed at both the transcriptional and post-transcriptional level, it is tempting to speculate that they promote senescence via two independent mechanisms. First, TE-derived nucleic acids can trigger innate sensing cascades that induce inflammation, IFN secretion and ultimately inflammaging. This proinflammatory cascade has been rigorously documented in the context of aging in murine and human cells [25,63,64]. Second, the retrotransposition of L1 and other mobile genetic elements directly induces DNA damage. Since several factors involved in DNA damage repair are downregulated in (pre)senescent cells (e.g. BRCA1, BRCA2, FANCA, FANCD2, etc.), these cells may not be able to repair the damage in a timely manner, which might ultimately lead to a permanent cell-cycle arrest.

## Conclusions

Our findings not only shed light on the differential expression of L1 modulators during the transition of cells to senescence, but also highlight the dual role that TE activation may play in senescence. On the one hand, our study supports a model, in which increased transcription of transposable elements, including L1 and Alu elements, is sensed and triggers IFN-mediated inflammaging. On the other hand, our analysis of Ribo-seq data revealed an increased translation of L1 elements that may enable L1 retrotransposition and ultimately induce genotoxic stress that further promotes transition to senescence. It will be important to decipher the relative contribution of these two mechanisms to senescence. Furthermore, future studies should aim at alleviating the detrimental effects of TEs on cellular senescence, organismal aging and aging-associated diseases.

## Declarations

### Ethics approval and consent to participate

Not Applicable.

### Consent for publication

Not Applicable.

### Availability of data and materials

All data generated or analyzed during this study are included in this published article and its supplementary information files. No new large datasets were generated or uploaded to repositories. The sources of publicly available datasets that were analyzed in the present study are provided in this article.

## Funding

DS was supported by the Heisenberg Program (grant ID: SA 2676/3-1) and CRC1506 (“Aging at Interfaces”) of the German Research Foundation (DFG).

### Authors’ contributions

A.A. conceptualized the study and performed most of the computational analyses. D.M. performed the qRT-PCR experiments. G.N. modified and provided BJ cells. D.S. and C.G. provided resources and acquired funding. A.A., D.S., M.S. and D.M. prepared the figures, and A.A. & M.S. wrote the initial draft of the manuscript. All authors reviewed and edited the manuscript.

## Acknowledgements

The authors would like to thank Isabell Haußmann for excellent technical assistance.

## Additional files

### Additional file 1. List of previously described L1 suppressors and their differential expression in replicative senescence

| Symbol | BaseMean | log2FoldChange<br>(Senescent vs.<br>non-senescent) | lfcSE | stat | pvalue | padj | PMIDs |
| --- | --- | --- | --- | --- | --- | --- | --- |
| ADAR | 13593.70544 | -0.1767522525 | 0.1036717693 | -1.704921732 | 0.08820902262 | 0.2020137145 | [52,53] |
| ADARB1 | 7923.720891 | -1.135034549 | 0.230490103 | -4.924439418 | 8.46E-07 | 5.84E-06 | [52] |
| APOBEC3A | 0.1771823279 | -1.285752751 | 3.470979875 | -0.3704293306 | 0.7110626231 | 0.8275012857 | [17,65–67] |
| APOBEC3B | 207.4065383 | -5.79319621 | 1.367403086 | -4.23664117 | 2.27E-05 | 1.28E-04 | [12–16] |
| APOBEC3C | 1361.025833 | 0.2410312535 | 0.2173211663 | 1.109101601 | 0.2673863542 | 0.4647505347 | [20] |
| APOBEC3F | 51.2280652 | 0.4755750593 | 0.3508502683 | 1.355492933 | 0.1752605586 | 0.3414929439 | [15,20] |
| APOBEC3G | 129.3980821 | 0.2909960237 | 0.6001933883 | 0.4848371032 | 6.28E-01 | 7.79E-01 | [19], [47] |
| BRCA1 | 1407.430926 | -4.512747793 | 0.1861865152 | -24.23778 | 8.90E-130 | 2.98E-127 | [68] |
| BRCA2 | 739.0177882 | -4.747624492 | 0.3692232744 | -12.85841067 | 7.71E-38 | 3.56E-36 | [68,69] |
| CDK12 | 2729.584381 | -0.3958855794 | 0.07516753684 | -5.266709487 | 1.39E-07 | 1.05E-06 | [70] |
| ERCC1 | 2590.195369 | -0.5402738228 | 0.1691851287 | -3.193388373 | 0.001406136906 | 0.005646594775 | [71–73] |
| FANCA | 859.0160672 | -4.567728483 | 0.2331840498 | -19.58851168 | 1.94E-85 | 3.27E-83 | [29,71] |
| FANCC | 576.3773425 | -1.160195003 | 0.1955136277 | -5.934087647 | 2.95E-09 | 2.69E-08 | [68,70,74] |
| FANCD2 | 1704.582755 | -4.578016147 | 0.3447290095 | -13.28004322 | 3.02E-40 | 1.51E-38 | [29,71] |
| FANCI | 2634.783325 | -3.87242164 | 0.1598756147 | -24.22146521 | 1.32E-129 | 4.39E-127 | [68–70,74] |
| MCM2 | 3296.152558 | -3.886187297 | 0.1677845159 | -23.16177555 | 1.11E-118 | 3.17E-116 | [75] |
| MEN1 | 630.8288085 | -0.4623118186 | 0.1703850649 | -2.713335344 | 6.66E-03 | 2.26E-02 | [68–70,74,76] |
| MORC2 | 1785.005624 | -0.4624012188 | 0.1333309595 | -3.468070886 | 5.24E-04 | 2.32E-03 | [36,77] |
| MOV10 | 2327.601271 | -0.4614617449 | 0.1198897 | -3.849052463 | 1.19E-04 | 5.92E-04 | [10,78] |
| PPHLN1 | 2641.473683 | -0.1631561449 | 0.1763207885 | -0.9253369741 | 3.55E-01 | 5.62E-01 | [79,80] |
| RNASEH2A | 1237.108977 | -3.591983858 | 0.133416489 | -26.92308789 | 1.18E-159 | 6.70E-157 | [81,82] |
| RNASEH2C | 684.8608255 | -0.699231256 | 0.2099357848 | -3.330691129 | 0.000866306685 | 0.003662943363 | [75,82–86] |
| SAFB | 2054.418039 | -1.028261863 | 0.1528576456 | -6.726924643 | 1.73E-11 | 1.95E-10 | [75,83–86] |
| SETDB1 | 1429.963425 | -0.3200143374 | 0.15809441 | -2.024197677 | 4.29E-02 | 1.13E-01 | [87–91] |
| SLFN5 | 5495.230925 | 1.451515478 | 0.1754247483 | 8.274291354 | 1.29E-16 | 2.17E-15 | [75,83–85] |
| SLX4 | 588.5899453 | -1.274654754 | 0.1839240204 | -6.930333247 | 4.20E-12 | 5.03E-11 | [29,71] |
| TASOR | 4660.656721 | 0.1989861321 | 0.12541377 | 1.586637034 | 1.13E-01 | 0.2441933186 | [36,77,79,92,<br>93] |
| TRIM5 | 1484.227229 | -0.4452832903 | 0.1658357538 | -2.685086178 | 0.007251110699 | 0.02432820151 | [75,83,84] |
| TRIM28 | 10838.28195 | -1.505814595 | 0.2386464441 | -6.309813672 | 2.79E-10 | 2.82E-09 | [75,83] |

### Additional file 2. List of previously described L1 suppressors and their differential expression in oncogene-induced senescence

| Symbol | BaseMean | log2FoldChange<br>(Senescent vs. non-senescent) | lfcSE | stat | pvalue | padj | PMID |
| --- | --- | --- | --- | --- | --- | --- | --- |
| ADAR | 8368.720992 | -0.0952368314 | 0.3484200601 | -0.2733391165 | 0.7845925491 | 0.9379457021 | [52,53] |
| ADARB1 | 784.1280316 | -1.794027878 | 0.4824583062 | -3.718513818 | 0.0002003983319 | 0.00490744613<br>2 | [52] |
| APOBEC3B | 310.1567351 | -6.752480232 | 0.9267465089 | -7.286221386 | 3.19E-13 | 4.08E-11 | [12–16] |
| APOBEC3C | 1932.589211 | 1.042916509 | 0.3744387161 | 2.785279577 | 5.35E-03 | 6.49E-02 | [20] |
| APOBEC3F | 96.05824889 | -1.044669274 | 0.5519057116 | -1.892840121 | 5.84E-02 | 0.3284414554 | [15] |
| APOBEC3G | 58.77648727 | -1.272260731 | 0.7374265801 | -1.725271051 | 0.08447863646 | 0.4083211076 | [15,19,47] |
| BRCA1 | 1153.869341 | -3.293584005 | 0.390952605 | -8.42450968 | 3.62E-17 | 7.06E-15 | [68] |
| BRCA2 | 638.2982856 | -4.015285616 | 0.4324966275 | -9.283969772 | 1.63E-20 | 4.09E-18 | [68,69] |
| CDK12 | 3317.522914 | 0.04627908005 | 0.3399569552 | 0.136132176 | 0.8917167915 | 0.9751049779 | [70] |
| ERCC1 | 2786.928375 | 1.293056276 | 0.3783880289 | 3.417275856 | 0.0006325115405 | 0.0126031845 | [71–73] |
| FANCA | 1279.213077 | -4.675477182 | 0.4690577733 | -9.967806629 | 2.11E-23 | 6.35E-21 | [29,71] |
| FANCC | 277.9076235 | -1.782393688 | 0.5618803281 | -3.172194504 | 0.001512916418 | 0.02486001012 | [68,70,74] |
| FANCD2 | 829.1968194 | -4.839139327 | 0.4218704277 | -11.4706768 | 1.85E-30 | 9.82E-28 | [29,71] |
| FANCI | 2143.908685 | -3.427379534 | 0.3631840345 | -9.43703249 | 3.83E-21 | 9.78E-19 | [68–70,74] |
| MCM2 | 3052.158556 | -3.440282643 | 0.4890269477 | -7.034955148 | 1.99E-12 | 2.42E-10 | [75] |
| MEN1 | 1266.918142 | -0.1952086275 | 0.3569306757 | -0.54690908 | 5.84E-01 | 9.14E-01 | [68–70,74,76] |
| MORC2 | 1392.51223 | -0.09762543581 | 0.34667252 | -0.2816070792 | 0.7782448114 | 0.9368147665 | [36,77] |
| MOV10 | 1887.955436 | 0.0634066136 | 0.3514920582 | 0.1803927347 | 0.8568442577 | 0.9660164808 | [10,78] |
| PPHLN1 | 1367.840596 | 0.4616055898 | 0.4073374611 | 1.133226462 | 2.57E-01 | 7.20E-01 | [79,80] |
| RNASEH2A | 1406.277496 | -3.624748877 | 0.4825712269 | -7.51132408 | 5.85E-14 | 8.21E-12 | [81,82] |
| RNASEH2C | 793.4435244 | 0.2589269923 | 0.3757228108 | 0.6891436583 | 4.91E-01 | 8.96E-01 | [75,82–86] |
| SAFB | 2466.688488 | -1.186810605 | 0.4269607716 | -2.779671303 | 0.005441394536 | 0.06559601113 | [75,83–86] |
| SETDB1 | 724.7476742 | -0.3170419275 | 0.371907219 | -0.8524758633 | 0.3939500299 | 0.8434003343 | [87–91] |
| SLFN5 | 4231.544048 | 1.199170999 | 0.712351448 | 1.683397994 | 0.09229807134 | 0.4295928367 | [75,83–85] |
| SLX4 | 896.9424992 | -0.3033578976 | 0.420107135 | -0.7220965137 | 0.4702351415 | 0.8843501764 | [29,71] |
| TASOR | 2524.530699 | -0.3932232306 | 0.4047780991 | -0.97145382 | 3.31E-01 | 7.98E-01 | [36,77,79,92,93] |
| **TRIM5** | 759.3398736 | -1.336946212 | 0.3767997659 | -3.54816094 | 0.0003879311041 | 0.00843996785<br>3 | [75,83,84] |
| **TRIM28** | 11634.48129 | -1.256029245 | 0.3900891791 | -3.219851542 | 0.001282569985 | 0.02184652445 | [75,83] |

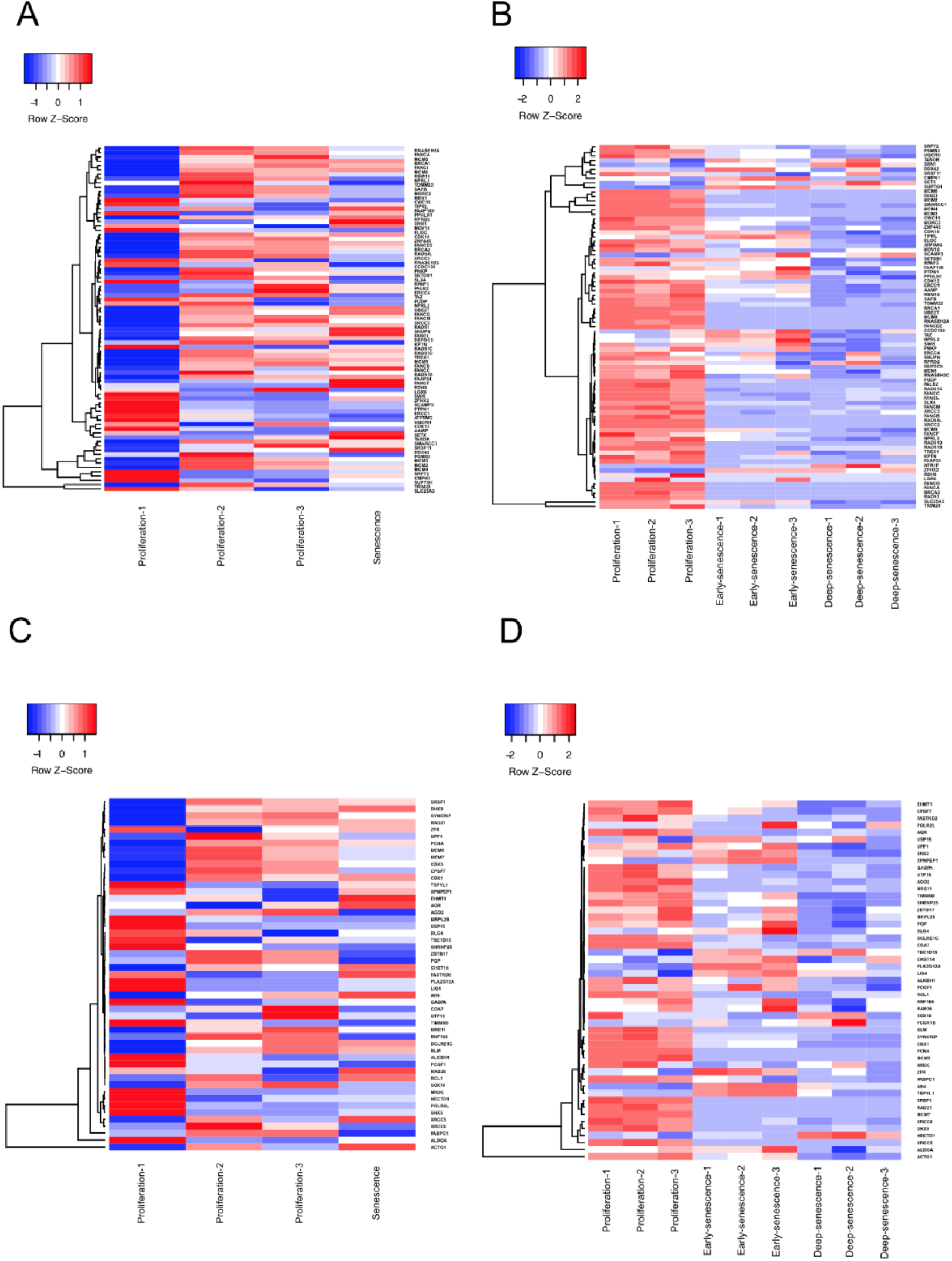

### **Additional file 3: Differential expression of L1 regulators in senescent vs. proliferating cells. (A-D)** Heatmaps illustrate the differential expression of (A, B) L1 suppressors and (C, D) L1 activators identified in a CRISPR/Cas screen by Liu and colleagues [36] in senescent vs. proliferating cells. Data were obtained from cells undergoing (A, C) oncogene-induced senescence [33] or (B, D) replicative senescence [25]

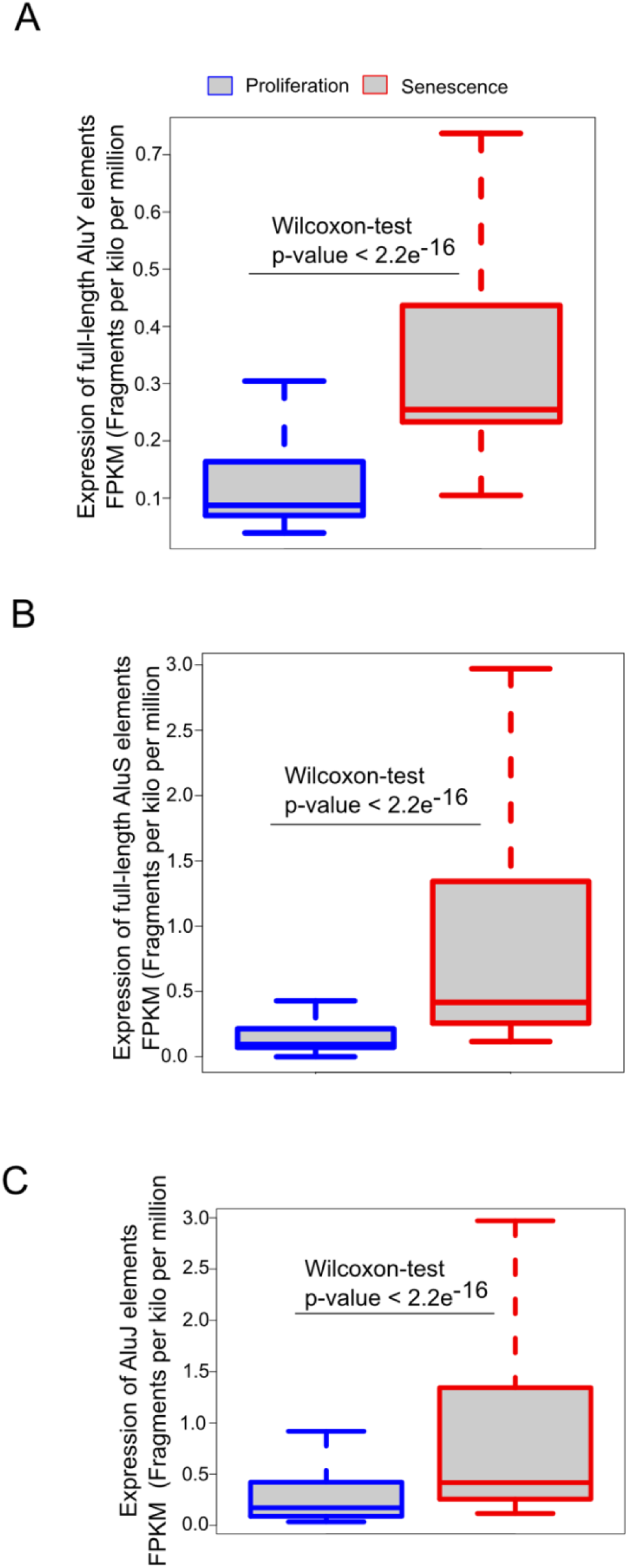

### **Additional file 4: Expression of Alu elements is significantly increased in senescent vs. proliferating cells. (A-C)** Expression of (A) AluY, (B) AluS and (C) AluJ elements in human BJ primary fibroblasts transitioning from a proliferating state to RAS^G12V^-induced senescence [33]. p-values were calculated using the Wilcoxon-Mann-Whitney test

## List of abbreviations

ADAR: Adenosine Deaminase Acting on RNA
Alu: Arthrobacter luteus
APOBEC3: Apolipoprotein B mRNA Editing Catalytic Polypeptide-like
bps: Base pairs
BRCA: Breast cancer gene
CPM: Counts per million
ES: Embryonic stem fibroblasts
EV: Empty vector
FANC: Fanconi anemia complementation group (FANC)
FPKM: Fragments Per Kilobase of transcript per Million mapped reads
GAPDH: Glyceraldehyde-3-Phosphate Dehydrogenase
GATK: Genome Analysis Toolkit
IFN: Interferon
L1: Long Interspersed Nuclear Elements-1
KAP1: KRAB-Associated Protein 1
KRAB-ZNF: Krüppel-Associated Box Zinc Finger Protein
log2fc: Log2 fold change
MCM2: Minichromosome Maintenance Complex Component 2
MOV10: Moloney leukemia virus 10
MPP8: M-Phase Phosphoprotein 8
ORF: Open reading frame
qRT-PCR: Quantitative Reverse Transcription Polymerase Chain Reaction
RAS^G12V^: Rat sarcoma mutant G12V
Ribo-seq: Ribosome Profiling Sequencing
RNA-seq: RNA sequencing
RNASEH2A: Ribonuclease H2 Subunit A
SASP: Senescence-associated secretory phenotype
SETDB1: Su(var)3-9, Enhancer-of-zeste, and Trithorax Domain Bifurcated Histone Lysine Methyltransferase 1
SINE: Short interspersed nuclear element
TASOR: Transcription Activation Suppressor
TE: Transposable element
TREX1: Three prime repair exonuclease 1
TRIM28: Tripartite motif-containing protein 28
UCSC: University of California, Santa Cruz
UHRF1: Ubiquitin like with PHD and ring finger domains 1

## References

1. Mills RE, Bennett EA, Iskow RC, Devine SE. Which transposable elements are active in the human genome? Trends Genet. 2007;23:183–91.

2. Kazazian HH Jr, Moran JV. Mobile DNA in Health and Disease. N Engl J Med. 2017;377:361–70.

3. Lander ES, Linton LM, Birren B, Nusbaum C, Zody MC, Baldwin J, et al. Initial sequencing and analysis of the human genome. Nature. 2001;409:860–921.

4. Salter JD, Bennett RP, Smith HC. The APOBEC Protein Family: United by Structure, Divergent in Function. Trends Biochem Sci. 2016;41:578–94.

5. Dewannieux M, Esnault C, Heidmann T. LINE-mediated retrotransposition of marked Alu sequences. Nat Genet. 2003;35:41–8.

6. Rowe HM, Jakobsson J, Mesnard D, Rougemont J, Reynard S, Aktas T, et al. KAP1 controls endogenous retroviruses in embryonic stem cells. Nature. 2010;463:237–40.

7. Seczynska M, Bloor S, Cuesta SM, Lehner PJ. Genome surveillance by HUSH-mediated silencing of intronless mobile elements. Nature. 2022;601:440–5.

8. Mathavarajah S, Dellaire G. LINE-1: an emerging initiator of cGAS-STING signalling and inflammation that is dysregulated in disease. Biochem Cell Biol. 2024;102:38–46.

9. Stetson DB, Ko JS, Heidmann T, Medzhitov R. Trex1 prevents cell-intrinsic initiation of autoimmunity. Cell. 2008;134:587–98.

10. Liu Q, Yi D, Ding J, Mao Y, Wang S, Ma L, et al. MOV10 recruits DCP2 to decap human LINE-1 RNA by forming large cytoplasmic granules with phase separation properties. EMBO Rep. 2023;24:e56512.

11. Arora R, Bodak M, Penouty L, Hackman C, Ciaudo C. Sequestration of LINE-1 in cytosolic aggregates by MOV10 restricts retrotransposition. EMBO Rep. 2022;23:e54458.

12. Wissing S, Montano M, Garcia-Perez JL, Moran JV, Greene WC. Endogenous APOBEC3B restricts LINE-1 retrotransposition in transformed cells and human embryonic stem cells. J Biol Chem. 2011;286:36427–37.

13. Lovsin N, Peterlin BM. APOBEC3 proteins inhibit LINE-1 retrotransposition in the absence of ORF1p binding. Ann N Y Acad Sci. 2009;1178:268–75.

14. Muckenfuss H, Hamdorf M, Held U, Perković M, Löwer J, Cichutek K, et al. APOBEC3 proteins inhibit human LINE-1 retrotransposition. J Biol Chem. 2006;281:22161–72.

15. Stenglein MD, Harris RS. APOBEC3B and APOBEC3F inhibit L1 retrotransposition by a DNA deamination-independent mechanism. J Biol Chem. 2006;281:16837–41.

16. Bogerd HP, Wiegand HL, Hulme AE, Garcia-Perez JL, O’Shea KS, Moran JV, et al. Cellular inhibitors of long interspersed element 1 and Alu retrotransposition. Proc Natl Acad Sci U S A. 2006;103:8780–5.

17. Richardson SR, Narvaiza I, Planegger RA, Weitzman MD, Moran JV. APOBEC3A deaminates transiently exposed single-strand DNA during LINE-1 retrotransposition. Elife. 2014;3:e02008.

18. Liang W, Xu J, Yuan W, Song X, Zhang J, Wei W, et al. APOBEC3DE Inhibits LINE-1 Retrotransposition by Interacting with ORF1p and Influencing LINE Reverse Transcriptase Activity. PLoS One. 2016;11:e0157220.

19. Koyama T, Arias JF, Iwabu Y, Yokoyama M, Fujita H, Sato H, et al. APOBEC3G oligomerization is associated with the inhibition of both Alu and LINE-1 retrotransposition. PLoS One. 2013;8:e84228.

20. Horn AV, Klawitter S, Held U, Berger A, Vasudevan AAJ, Bock A, et al. Human LINE-1 restriction by APOBEC3C is deaminase independent and mediated by an ORF1p interaction that affects LINE reverse transcriptase activity. Nucleic Acids Res. 2014;42:396–416.

21. Prior S, Miousse IR, Nzabarushimana E, Pathak R, Skinner C, Kutanzi KR, et al. Densely ionizing radiation affects DNA methylation of selective LINE-1 elements. Environ Res. 2016;150:470–81.

22. Singh A, Rappolee DA, Ruden DM. Epigenetic Reprogramming in and : From Fertilization to Primordial Germ Cell Development. Cells [Internet]. 2023;12. Available from: 10.3390/cells12141874

23. Min B, Jeon K, Park JS, Kang Y-K. Demethylation and derepression of genomic retroelements in the skeletal muscles of aged mice. Aging Cell. 2019;18:e13042.

24. Ndhlovu LC, Bendall ML, Dwaraka V, Pang APS, Dopkins N, Carreras N, et al. Retro-age: A unique epigenetic biomarker of aging captured by DNA methylation states of retroelements. Aging Cell. 2024;23:e14288.

25. De Cecco M, Ito T, Petrashen AP, Elias AE, Skvir NJ, Criscione SW, et al. L1 drives IFN in senescent cells and promotes age-associated inflammation. Nature. 2019;566:73–8.

26. Simon M, Van Meter M, Ablaeva J, Ke Z, Gonzalez RS, Taguchi T, et al. LINE1 Derepression in Aged Wild-Type and SIRT6-Deficient Mice Drives Inflammation. Cell Metab. 2019;29:871–85.e5.

27. Zarneshan SN, Fakhri S, Bachtel G, Bishayee A. Exploiting pivotal mechanisms behind the senescence-like cell cycle arrest in cancer. Adv Protein Chem Struct Biol. 2023;135:1– 19.

28. Kumari R, Jat P. Mechanisms of Cellular Senescence: Cell Cycle Arrest and Senescence Associated Secretory Phenotype. Front Cell Dev Biol. 2021;9:645593.

29. Brégnard C, Guerra J, Déjardin S, Passalacqua F, Benkirane M, Laguette N. Upregulated LINE-1 Activity in the Fanconi Anemia Cancer Susceptibility Syndrome Leads to Spontaneous Pro-inflammatory Cytokine Production. EBioMedicine. 2016;8:184–94.

30. da Silva PFL, Ogrodnik M, Kucheryavenko O, Glibert J, Miwa S, Cameron K, et al. The bystander effect contributes to the accumulation of senescent cells in vivo. Aging Cell. 2019;18:e12848.

31. Ramini D, Latini S, Giuliani A, Matacchione G, Sabbatinelli J, Mensà E, et al. Replicative Senescence-Associated LINE1 Methylation and LINE1-Alu Expression Levels in Human Endothelial Cells. Cells [Internet]. 2022;11. Available from: 10.3390/cells11233799

32. Tang H, Yang J, Xu J, Zhang W, Geng A, Jiang Y, et al. The transcription factor PAX5 activates human LINE1 retrotransposons to induce cellular senescence. EMBO Rep. 2024;25:3263–75.

33. Loayza-Puch F, Drost J, Rooijers K, Lopes R, Elkon R, Agami R. p53 induces transcriptional and translational programs to suppress cell proliferation and growth. Genome Biol. 2013;14:R32.

34. Dobin A, Davis CA, Schlesinger F, Drenkow J, Zaleski C, Jha S, et al. STAR: ultrafast universal RNA-seq aligner. Bioinformatics. 2013;29:15–21.

35. Liao Y, Smyth GK, Shi W. featureCounts: an efficient general purpose program for assigning sequence reads to genomic features. Bioinformatics. 2014;30:923–30.

36. Liu N, Lee CH, Swigut T, Grow E, Gu B, Bassik MC, et al. Selective silencing of euchromatic L1s revealed by genome-wide screens for L1 regulators. Nature. 2018;553:228– 32.

37. Langmead B, Salzberg SL. Fast gapped-read alignment with Bowtie 2. Nat Methods. 2012;9:357–9.

38. Trapnell C, Williams BA, Pertea G, Mortazavi A, Kwan G, van Baren MJ, et al. Transcript assembly and quantification by RNA-Seq reveals unannotated transcripts and isoform switching during cell differentiation. Nat Biotechnol. 2010;28:511–5.

39. Kent WJ, Sugnet CW, Furey TS, Roskin KM, Pringle TH, Zahler AM, et al. The human genome browser at UCSC. Genome Res. 2002;12:996–1006.

40. McKenna A, Hanna M, Banks E, Sivachenko A, Cibulskis K, Kernytsky A, et al. The Genome Analysis Toolkit: a MapReduce framework for analyzing next-generation DNA sequencing data. Genome Res. 2010;20:1297–303.

41. Piskol R, Ramaswami G, Li JB. Reliable identification of genomic variants from RNA-seq data. Am J Hum Genet. 2013;93:641–51.

42. Quinlan AR, Hall IM. BEDTools: a flexible suite of utilities for comparing genomic features. Bioinformatics. 2010;26:841–2.

43. Kim D, Pertea G, Trapnell C, Pimentel H, Kelley R, Salzberg SL. TopHat2: accurate alignment of transcriptomes in the presence of insertions, deletions and gene fusions. Genome Biol. 2013;14:R36.

44. Ramírez F, Dündar F, Diehl S, Grüning BA, Manke T. deepTools: a flexible platform for exploring deep-sequencing data. Nucleic Acids Res. 2014;42:W187–91.

45. Turelli P, Castro-Diaz N, Marzetta F, Kapopoulou A, Raclot C, Duc J, et al. Interplay of TRIM28 and DNA methylation in controlling human endogenous retroelements. Genome Res. 2014;24:1260–70.

46. Orecchini E, Frassinelli L, Galardi S, Ciafrè SA, Michienzi A. Post-transcriptional regulation of LINE-1 retrotransposition by AID/APOBEC and ADAR deaminases. Chromosome Res. 2018;26:45–59.

47. Khatua AK, Taylor HE, Hildreth JEK, Popik W. Inhibition of LINE-1 and Alu retrotransposition by exosomes encapsidating APOBEC3G and APOBEC3F. Virology. 2010;400:68–75.

48. Jurka J, Smith T. A fundamental division in the Alu family of repeated sequences. Proc Natl Acad Sci U S A. 1988;85:4775–8.

49. Batzer MA, Deininger PL, Hellmann-Blumberg U, Jurka J, Labuda D, Rubin CM, et al. Standardized nomenclature for Alu repeats. J Mol Evol. 1996;42:3–6.

50. Sanchez A, Ortega P, Sakhtemani R, Manjunath L, Oh S, Bournique E, et al. Mesoscale DNA features impact APOBEC3A and APOBEC3B deaminase activity and shape tumor mutational landscapes. Nat Commun. 2024;15:2370.

51. Savva YA, Rieder LE, Reenan RA. The ADAR protein family. Genome Biol. 2012;13:252.

52. Frassinelli L, Orecchini E, Al-Wardat S, Tripodi M, Mancone C, Doria M, et al. The RNA editing enzyme ADAR2 restricts L1 mobility. RNA Biol. 2021;18:75–87.

53. Orecchini E, Doria M, Antonioni A, Galardi S, Ciafrè SA, Frassinelli L, et al. ADAR1 restricts LINE-1 retrotransposition. Nucleic Acids Res. 2017;45:155–68.

54. Choudhary S, Li W, D Smith A. Accurate detection of short and long active ORFs using Ribo-seq data. Bioinformatics. 2020;36:2053–9.

55. Doehle BP, Schäfer A, Cullen BR. Human APOBEC3B is a potent inhibitor of HIV-1 infectivity and is resistant to HIV-1 Vif. Virology. 2005;339:281–8.

56. Kinomoto M, Kanno T, Shimura M, Ishizaka Y, Kojima A, Kurata T, et al. All APOBEC3 family proteins differentially inhibit LINE-1 retrotransposition. Nucleic Acids Res. 2007;35:2955–64.

57. Covino DA, Gauzzi MC, Fantuzzi L. Understanding the regulation of APOBEC3 expression: Current evidence and much to learn. J Leukoc Biol. 2018;103:433–44.

58. Rosser JM, An W. L1 expression and regulation in humans and rodents. Front Biosci (Elite Ed). 2012;4:2203–25.

59. Stenz L. The L1-dependant and Pol III transcribed Alu retrotransposon, from its discovery to innate immunity. Mol Biol Rep. 2021;48:2775–89.

60. Callinan PA, Wang J, Herke SW, Garber RK, Liang P, Batzer MA. Alu retrotransposition-mediated deletion. J Mol Biol. 2005;348:791–800.

61. Wang K, Li Y, Dai C, Wang K, Yu J, Tan Y, et al. Characterization of the relationship between APOBEC3B deletion and ACE Alu insertion. PLoS One. 2013;8:e64809.

62. Unger MA, Nathanson KL, Calzone K, Antin-Ozerkis D, Shih HA, Martin AM, et al. Screening for genomic rearrangements in families with breast and ovarian cancer identifies BRCA1 mutations previously missed by conformation-sensitive gel electrophoresis or sequencing. Am J Hum Genet. 2000;67:841–50.

63. Reid Cahn A, Bhardwaj N, Vabret N. Dark genome, bright ideas: Recent approaches to harness transposable elements in immunotherapies. Cancer Cell. 2022;40:792–7.

64. Gazquez-Gutierrez A, Witteveldt J, R Heras S, Macias S. Sensing of transposable elements by the antiviral innate immune system. RNA. 2021;27:735–52.

65. Bulliard Y, Narvaiza I, Bertero A, Peddi S, Röhrig UF, Ortiz M, et al. Structure-function analyses point to a polynucleotide-accommodating groove essential for APOBEC3A restriction activities. J Virol. 2011;85:1765–76.

66. Miyoshi T, Makino T, Moran JV. Poly(ADP-Ribose) Polymerase 2 Recruits Replication Protein A to Sites of LINE-1 Integration to Facilitate Retrotransposition. Mol Cell. 2019;75:1286–98.e12.

67. Mitra M, Hercík K, Byeon I-JL, Ahn J, Hill S, Hinchee-Rodriguez K, et al. Structural determinants of human APOBEC3A enzymatic and nucleic acid binding properties. Nucleic Acids Res. 2014;42:1095–110.

68. Carugno M, Maggioni C, Crespi E, Bonzini M, Cuocina S, Dioni L, et al. Night Shift Work, DNA Methylation and Telomere Length: An Investigation on Hospital Female Nurses. Int J Environ Res Public Health [Internet]. 2019;16. Available from: 10.3390/ijerph16132292

69. Ketola K, Kaljunen H, Taavitsainen S, Kaarijärvi R, Järvelä E, Rodríguez-Martín B, et al. Subclone Eradication Analysis Identifies Targets for Enhanced Cancer Therapy and Reveals L1 Retrotransposition as a Dynamic Source of Cancer Heterogeneity. Cancer Res. 2021;81:4901–9.

70. Bamberger C, Pankow S, Yates JR 3rd. SMG1 and CDK12 Link ΔNp63α Phosphorylation to RNA Surveillance in Keratinocytes. J Proteome Res. 2021;20:5347–58.

71. Bona N, Crossan GP. Fanconi anemia DNA crosslink repair factors protect against LINE-1 retrotransposition during mouse development. Nat Struct Mol Biol. 2023;30:1434–45.

72. Servant G, Streva VA, Derbes RS, Wijetunge MI, Neeland M, White TB, et al. The Nucleotide Excision Repair Pathway Limits L1 Retrotransposition. Genetics. 2017;205:139– 53.

73. Gasior SL, Roy-Engel AM, Deininger PL. ERCC1/XPF limits L1 retrotransposition. DNA Repair (Amst). 2008;7:983–9.

74. Tristan-Ramos P, Morell S, Sanchez L, Toledo B, Garcia-Perez JL, Heras SR. sRNA/L1 retrotransposition: using siRNAs and miRNAs to expand the applications of the cell culture-based LINE-1 retrotransposition assay. Philos Trans R Soc Lond B Biol Sci. 2020;375:20190346.

75. Xu B, Li X, Zhang S, Lian M, Huang W, Zhang Y, et al. Pan cancer characterization of genes whose expression has been associated with LINE-1 antisense promoter activity. Mob DNA. 2023;14:13.

76. Simbolo M, Bilotta M, Mafficini A, Luchini C, Furlan D, Inzani F, et al. Gene Expression Profiling of Pancreas Neuroendocrine Tumors with Different Ki67-Based Grades. Cancers (Basel) [Internet]. 2021;13. Available from: 10.3390/cancers13092054

77. Pandiloski N, Horváth V, Karlsson O, Koutounidou S, Dorazehi F, Christoforidou G, et al. DNA methylation governs the sensitivity of repeats to restriction by the HUSH-MORC2 corepressor. Nat Commun. 2024;15:7534.

78. Goodier JL, Wan H, Soares AO, Sanchez L, Selser JM, Pereira GC, et al. ZCCHC3 is a stress granule zinc knuckle protein that strongly suppresses LINE-1 retrotransposition. PLoS Genet. 2023;19:e1010795.

79. Jensvold ZD, Christenson AE, Flood JR, Lewis PW. Interplay between Two Paralogous Human Silencing Hub (HuSH) Complexes in Regulating LINE-1 Element Silencing [Internet]. bioRxiv. 2024. Available from: 10.1101/2023.12.28.573526

80. Douse CH, Tchasovnikarova IA, Timms RT, Protasio AV, Seczynska M, Prigozhin DM, et al. TASOR is a pseudo-PARP that directs HUSH complex assembly and epigenetic transposon control. Nat Commun. 2020;11:4940.

81. Choi J, Hwang S-Y, Ahn K. Interplay between RNASEH2 and MOV10 controls LINE-1 retrotransposition. Nucleic Acids Res. 2018;46:1912–26.

82. Pokatayev V, Hasin N, Chon H, Cerritelli SM, Sakhuja K, Ward JM, et al. RNase H2 catalytic core Aicardi-Goutières syndrome-related mutant invokes cGAS-STING innate immune-sensing pathway in mice. J Exp Med. 2016;213:329–36.

83. Robbez-Masson L, Tie CHC, Conde L, Tunbak H, Husovsky C, Tchasovnikarova IA, et al. The HUSH complex cooperates with TRIM28 to repress young retrotransposons and new genes. Genome Res. 2018;28:836–45.

84. Volkmann B, Wittmann S, Lagisquet J, Deutschmann J, Eissmann K, Ross JJ, et al. Human TRIM5α senses and restricts LINE-1 elements. Proc Natl Acad Sci U S A. 2020;117:17965–76.

85. Ding J, Wang S, Liu Q, Duan Y, Cheng T, Ye Z, et al. Schlafen-5 inhibits LINE-1 retrotransposition. iScience. 2023;26:107968.

86. Xiong F, Wang R, Lee J-H, Li S, Chen S-F, Liao Z, et al. RNA mA modification orchestrates a LINE-1-host interaction that facilitates retrotransposition and contributes to long gene vulnerability. Cell Res. 2021;31:861–85.

87. Fernandes LP, Enriquez-Gasca R, Gould PA, Holt JH, Conde L, Ecco G, et al. A satellite DNA array barcodes chromosome 7 and regulates totipotency via ZFP819. Sci Adv. 2022;8:eabp8085.

88. Müller I, Moroni AS, Shlyueva D, Sahadevan S, Schoof EM, Radzisheuskaya A, et al. MPP8 is essential for sustaining self-renewal of ground-state pluripotent stem cells. Nat Commun. 2021;12:3034.

89. Healton SE, Pinto HD, Mishra LN, Hamilton GA, Wheat JC, Swist-Rosowska K, et al. H1 linker histones silence repetitive elements by promoting both histone H3K9 methylation and chromatin compaction. Proc Natl Acad Sci U S A. 2020;117:14251–8.

90. Fukuda K, Shinkai Y. SETDB1-Mediated Silencing of Retroelements. Viruses [Internet]. 2020;12. Available from: 10.3390/v12060596

91. Jurkowska RZ, Qin S, Kungulovski G, Tempel W, Liu Y, Bashtrykov P, et al. H3K14ac is linked to methylation of H3K9 by the triple Tudor domain of SETDB1. Nat Commun. 2017;8:2057.

92. Danac JMC, Matthews RE, Gungi A, Qin C, Parsons H, Antrobus R, et al. Competition between two HUSH complexes orchestrates the immune response to retroelement invasion. Mol Cell. 2024;84:2870–81.e5.

93. Li Z, Duan S, Hua X, Xu X, Li Y, Menolfi D, et al. Asymmetric distribution of parental H3K9me3 in S phase silences L1 elements. Nature. 2023;623:643–51.

94. Meena, Jitendra, K Lenhard Rudolph, and Cagatay Günes. 2015. “Telomere Dysfunction, Chromosomal Instability and Cancer.” Recent Results in Cancer Research. Fortschritte Der Krebsforschung. Progres Dans Les Recherches Sur Le Cancer 200: 61–79.

95. Meessen, Sabine, Gregoire Najjar, Anca Azoitei, Sebastian Iben, Christian Bolenz, and Cagatay Günes. 2022. “A Comparative Assessment of Replication Stress Markers in the Context of Telomerase.” Cancers 14 (9). 10.3390/cancers14092205.

